# Surveying the contribution of rare variants to the genetic architecture of human disease through exome sequencing of 177,882 UK Biobank participants

**DOI:** 10.1101/2020.12.13.422582

**Authors:** Quanli Wang, Ryan S. Dhindsa, Keren Carss, Andrew Harper, Abhishek Nag, Ioanna Tachmazidou, Dimitrios Vitsios, Sri VV Deevi, Alex Mackay, Daniel Muthas, Michael Hühn, Susan Monkley, Henric Olsson, Sebastian Wasilewski, Katherine R. Smith, Ruth March, Adam Platt, Carolina Haefliger, Slavé Petrovski, AstraZeneca Genomics Initiative

## Abstract

The UK Biobank (UKB) represents an unprecedented population-based study of 502,543 participants with detailed phenotypic data and linkage to medical records. While the release of genotyping array data for this cohort has bolstered genomic discovery for common variants, the contribution of rare variants to this broad phenotype collection remains relatively unknown. Here, we use exome sequencing data from 177,882 UKB participants to evaluate the association between rare protein-coding variants with 10,533 binary and 1,419 quantitative phenotypes. We performed both a variant-level phenome-wide association study (PheWAS) and a gene-level collapsing analysis-based PheWAS tailored to detecting the aggregate contribution of rare variants. The latter revealed 911 statistically significant gene-phenotype relationships, with a median odds ratio of 15.7 for binary traits. Among the binary trait associations identified using collapsing analysis, 83% were undetectable using single variant association tests, emphasizing the power of collapsing analysis to detect signal in the setting of high allelic heterogeneity. As a whole, these genotype-phenotype associations were significantly enriched for loss-of-function mediated traits and currently approved drug targets. Using these results, we summarise the contribution of rare variants to common diseases in the context of the UKB phenome and provide an example of how novel gene-phenotype associations can aid in therapeutic target prioritisation.

## Introduction

Identifying genetic variants that contribute to human disease has facilitated the development of highly efficacious and safe therapeutics.^1–3^ It is now recognised that human genetic evidence supporting a drug target increases the likelihood of approval by at least two-fold.^4,5^ The UK Biobank (UKB), which integrates genetic data with phenotypic data linked to electronic health records for approximately 500,000 individuals, has provided a medical research resource of unprecedented scale. The release of genotyping array data for this cohort has ushered in a new era of genomic discovery through genome-wide association studies (GWAS) focused on common variants.^6,7^ However, there are two factors that make it challenging to directly translate such genetic associations into potential therapeutic opportunities. First, most common variants only have a modest effect on phenotype compared to rare variants. Second, it is difficult to precisely map complex disease associations to causal genes due to linkage disequilibrium and the fact that the implicated variants are often non-coding, which complicates efforts to translate such genetic associations into potential therapeutic opportunities.

It is well-recognised that rare functional variants tend to have larger phenotypic effects,^8^ enhancing their value for gleaning biological insight into disease. However, due to a lack of large-scale clinico-genomic datasets, the contribution of rare variants has until recently only been assessed for a subset of complex traits. The genome Aggregation Database (gnomAD), which includes exome and genome sequencing data of 141,456 individuals, constitutes the largest publicly available next-generation sequencing resource to date.^9,10^ While this resource has undeniably transformed our ability to interpret rare variants and characterise disease-associated genes, it is unsuited for the systematic assessment of the contribution of rare variation to human disease because of a lack of linked phenotypic data. Recent analyses of smaller sequenced collections with linked phenotypic data, including the first release of exome sequencing data for 50,000 UKB participants, have indicated an important role of rare variation in complex disease, highlighting a need for larger sample sizes to better understand this contribution.^11,12^

In this study, we analyse exome sequence data from 177,882 unrelated European ancestry UKB participants to evaluate the association between protein-coding variants with 10,533 binary and 1,419 quantitative phenotypes. First, we present the diversity of phenotypes and sequence variation captured in this cohort. We then perform variant- and gene-level association tests to identify protein-coding genetic risk factors across the allele frequency spectrum for thousands of clinical and quantitative traits. To illustrate the value of this approach, we perform molecular studies to interrogate the biological effects of a novel association between hemicentin 1 (*HMCN1)* expression and lung function. These analyses collectively provide a comprehensive catalogue of the contribution from protein-coding variation to the genetic architecture of a broad range of complex human diseases and biomarkers.

## Results

### Clinical and demographic characterization of the cohort

We analysed exome sequence data from 200,593 UKB participants, totalling more than 665 terabytes of raw sequencing data, which was processed through a standardised, cloud-based bioinformatics pipeline **(methods)**. We performed stringent quality control to remove samples with low sequencing quality, and we restricted cohort analyses to unrelated index samples (**methods**).

Participant data includes periodically updated health records, self-reported survey information, linkage to death and cancer registries, quantitative biomarkers, imaging data and other phenotypic endpoints.^7^ Due to the variable categorisation modes, scaling, and follow-up responses inherent to this data, we created a modified version of the previously introduced PHESANT package (see methods),^13^ in order to systematically harmonise phenotypes. In total, we considered 10,533 binary traits and 1,419 quantitative traits, which we categorised into 22 distinct ICD-10-based chapters (**Fig. 1a, b, Supplementary Table 1**). Due to the correlative structure among the binary traits, we also introduced a union mapping approach, in which we attempted to group similar phenotypes together (**methods, Supplementary Table 1**). Therefore 4,817 of the 10,533 binary phenotypes are union phenotypes.

**Figure 1.**
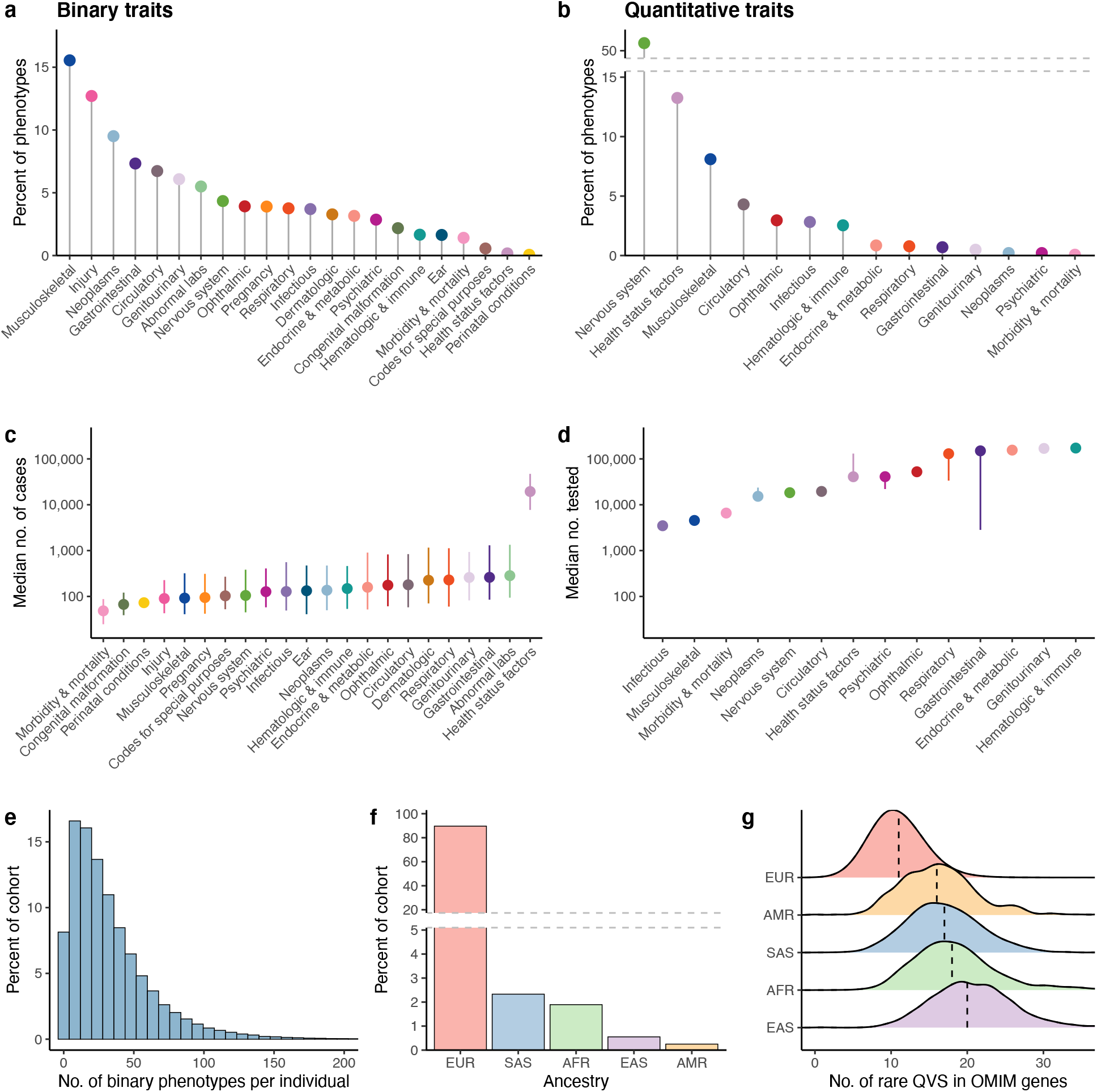
Phenotypic and demographic diversity of the sequenced UK Biobank cohort. **(a)** The percentage of binary union traits assessed in the cohort that correspond to the chapter. **(b)** The percentage of quantitative traits assessed in the cohort per chapter. **(c)** The median number of cases of European ancestry per binary union phenotype stratified by chapter. **(d)** The median number of participants of European ancestry tested for quantitative traits stratified by chapter. **(e)** Histogram depicting the number of binary union phenotypes per patient. The x-axis was capped at 200 for visual clarity. **(f)** The distribution of represented genetic ancestries in the sequenced cohort. EUR = European, SAS = South Asian, AFR = African, EAS = East Asian, AMR = American. **(g)** The distribution of the number of ultra-rare (MAF < 0.005%) qualifying variants (QVs) in OMIM-derived Mendelian disease genes per ancestral group. Error bars in (c, d) represent the interquartile range.

The average age at recruitment for sequenced individuals was 56.7 years and 55% of the sequenced cohort were female. All variant- and gene-level association tests were performed in individuals of European genetic ancestry **(methods).** The median number of cases per binary union phenotype is 132 (interquartile range: 49-527) (**Fig. 1c, d**) and the median number of binary union-mapped traits per participant is 25 (interquartile range: 12-45) of the possible 4,817 (**Fig. 1e**).

As we sequence more individuals from a given population, our resolution to evaluate variants across the allele frequency spectrum increases. When one considers ultra-rare (MAF<0.005%) non-synonymous variants among OMIM disease-associated genes, individuals of European ancestry (**Fig. 1f**) have a substantially higher number of such candidate variants (**Fig. 1g**). This indicates that the dearth of available sequencing data in non-European ancestries reduces our ability to effectively detect pathogenic variants in those ancestries, as has also previously been observed.^14^ It is therefore crucial that global biobanks that represent individuals of diverse genetic ancestries work towards providing reference data similar to the UKB for the benefit of the international medical community.

### Variant-level exome-wide association studies

Exome sequencing allows us to test for the association between phenotypes and individual protein-coding variants across the allele frequency spectrum. The tendency of rare variants to have a larger effect on disease risk underscores the value of this dataset in assessing their contribution on a phenome-wide scale. Protein-truncating variants (PTVs), which are predicted to shorten the coding sequence of genes, constitute one class of variation that has revealed much about human biology and disease mechanisms.^10,15^ Furthermore, these variants hold promise for drug discovery since identifying PTVs that protect against human disease could provide direct human validation of potential therapeutic targets.^3,16^ By surveying sequence data from 191,022 UKB participants of any ancestry, we observed that 96% of human protein-coding genes had at least one heterozygous putative PTV carrier, 40% had at least one putatively compound heterozygous or homozygous PTV carrier, and 18% of the 18,741 studied genes had at least one homozygous/hemizygous PTV carrier (**Fig. 2a**). Among the 191,022 UKB participants, only 899 genes (4.7%) harboured a PTV with a MAF > 0.5%, a threshold comfortably captured via microarray technology (**Fig. 2a**), further emphasizing the power of exome-sequencing in studying this crucial class of genetic variation.

**Figure 2.**
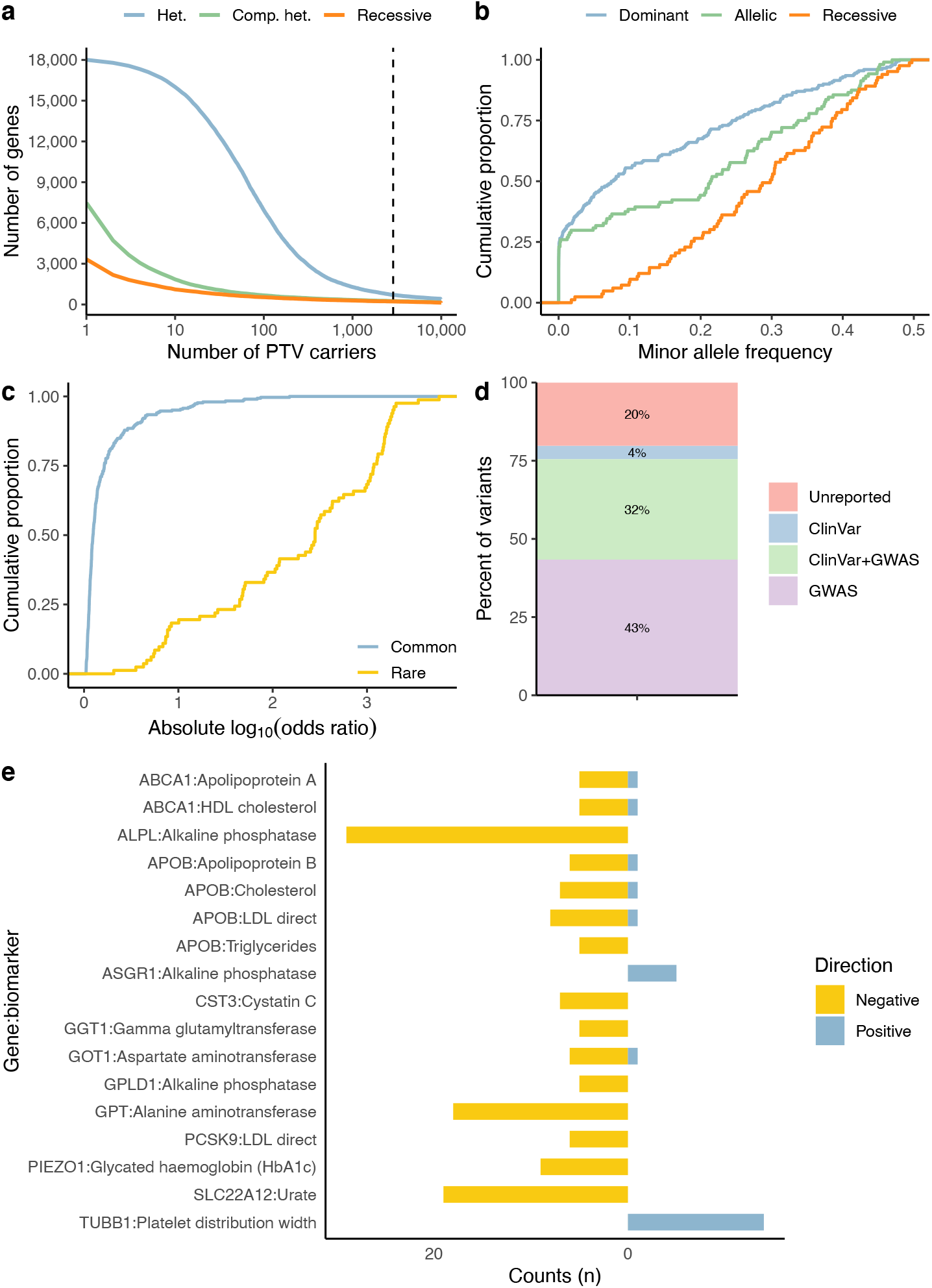
Summary of variant-level exome-wide association study results. **(a)** The number of genes (y-axis) with at least *N* protein-truncating variant (PTV) carriers (x-axis) in the cohort. The dotted line corresponds to (MAF > 0.5%, i.e. 2,873 carriers), the number of carriers typically reliably detected via array technologies. Colours correspond to heterozygous (Het), putative compound heterozygous (comp. het), and homozygous/hemizygous carriers (recessive). **(b)** The MAF distribution of genome-wide significant ExWAS variants across all binary phenotypes assessed. **(c)** The distribution of effect sizes for common (MAF≥0.5%) compared to rare (MAF < 0.5%) significant ExWAS variants. In plots b and c, we display the variants with the largest effect sizes achieved per gene. **(d)** Percentage of ExWAS study-wide significant PTVs or missense variants that were previously reported in ClinVar or the GWAS catalogue (including GWAS Catalogue variants within 50kbp flanking sequence either side of the index variant). **(e)** Distribution of the directions of effect for the rare non-synonymous variant associations for the significant gene-phenotype relationships. Only quantitative phenotypes with at least five significant non-synonymous variant associations (p < 1 × 10^−8^) in a given gene were considered.

We performed an exome-wide association study (ExWAS) across all phenotypes to test for genetic associations with variants that were observed among at least 6 of the 177,882 European ancestry participants (equivalent to a MAF lower limit of 0.001%) **(methods)**. Adopting a p-value threshold of *P*<1×10^−8^ (**methods**), we identified a total of 2,719 distinct genotype-phenotype associations for binary traits and 28,465 distinct associations for quantitative traits, outside of the MHC region as defined here by chr6:25Mbp-35Mbp (**Table 1 and Supplementary Table 2**). The variant with the lowest allele frequency that achieved study-wide significance was a Uromodulin (*UMOD)* frameshift variant associated with Chronic kidney disease, stage 5 (cohort MAF of 0.0019%) (**Table 1**). Overall, we found that many of the significant ExWAS signals were explained by variants with a MAF<0.5% (**Fig 2b**). For the dominant model in particular, variants with a MAF<0.5% account for 27.5% of all statistically significant variant-level associations.

**Table 1.** Top PTV ExWAS signals. Restricting to ExWAS protein-truncating variant signals that achieved a p-value < 1×10^−8^ in at least one of the three studied genetic models for binary traits (allelic, dominant or recessive). Not including variants within the MHC region, defined here as chr6:25Mbp– 35Mbp. For table brevity only the phenotype with the greatest effect size is presented for each unique variant. Shaded lines = control MAF < 0.5%. ClinSigSimple data is provided by ClinVar where 0 = no current value of Likely pathogenic or Pathogenic and 1= at least one current record submitted with an interpretation of Likely pathogenic or Pathogenic (independent of whether that record includes assertion criteria and evidence).

| Gene | Variant | Most Damaging | Model | Phenotype | p-value | OR | Case MAF | Ctrl MAF | ClinVar ClinSig Simple | OMIM |
| --- | --- | --- | --- | --- | --- | --- | --- | --- | --- | --- |
| *ABO* | 9-133257521-T-TC | splice acceptor | dom | 41202#I802#I80.2 Phlebitis and thrombophlebitis of other deep vessels of lower extremities | 2.82E-22 | 1.8 | 0.432 | 0.3347 | NA | [Blood group; ABO system] |
| *HBB* | 11-5226774-G-A | stop gained | dom | Union#D56#D56 Thalassaemia | 1.05E-18 | 3497.8 | 0.0448 | 1.4E-05 | 1 | Delta-beta thalassemia |
| *MYOC* | 1-171636338-G-A | stop gained | dom | Union#H409#H40.9 Glaucoma\| unspecified | 1.52E-17 | 6.2 | 0.0078 | 0.0013 | 1 | Glaucoma 1A; primary open angle |
| *COL4A4* | 2-227052367-G-C | stop gained | dom | Union#R31#R31 Unspecified haematuria | 2.15E-16 | 5.6 | 0.0026 | 4.7E-04 | 1 | Hematuria; familial benign |
| *FLG* | 1-152312600-CACTG-C | frameshift | rec | 20002#1452#eczema\|dermatitis | 1.75E-14 | 9.7 | 0.0459 | 0.0235 | 1 | Ichthyosis vulgaris |
|  | 1-152313385-G-A | stop gained | rec | 20002#1452#eczema\|dermatitis | 6.27E-14 | 10.5 | 0.0454 | 0.0229 | 1 | Ichthyosis vulgaris |
| *UMOD* | 16-20349020-CA-C | frameshift | dom | Union#N185#N18.5 Chronic kidney disease\| stage 5 | 7.14E-13 | 1933.8 | 0.0074 | 3.9E-06 | NA | Glomerulocystic kidney disease with hyperuricemia and isosthenuria |
|  | 16-20349017-G-GA | frameshift | dom | Union#D638#D63.8 Anaemia in other chronic diseases classified elsewhere | 6.42E-11 | 1403.5 | 0.0088 | 6.4E-06 | NA | Glomerulocystic kidney disease with hyperuricemia and isosthenuria |
|  | 16-20349015-C-A | stop gained | dom | 41202#N185#N18.5 Chronic kidney disease\| stage 5 | 2.16E-10 | 1036.6 | 0.0097 | 9.5E-06 | 0 | Glomerulocystic kidney disease with hyperuricemia and isosthenuria |
| *CHEK2* | 22-28695868-AG-A | frameshift | dom | 20001#1002#breast cancer | 4.90E-12 | 3.5 | 0.0059 | 0.0017 | 1 | {Breast and colorectal cancer; susceptibility to} |
| *MROH2A* | 2-233823665-G-A | splice donor | dom | Source of report of E80 (disorders of porphyrin and bilirubin metabolism) | 1.25E-11 | 1.8 | 0.2393 | 0.1577 | NA | NA |
| *MUC1* | 1-155192276-C-T | splice acceptor | allele | Union#K317#K31.7 Polyp of stomach and duodenum | 2.30E-10 | 1.2 | 0.4262 | 0.463 | NA | Medullary cystic kidney disease |
| *FES* | 15-90885291-CT-C | frameshift | rec | 6150#4#Vascular/heart problems diagnosed by doctor\| High Blood Pressure | 2.37E-10 | 1.1 | 0.3375 | 0.3222 | NA | NA |
| *TSPAN10* | 17-81647906-T-TTAAC | frameshift | rec | Source of report of H26 (other cataract) | 2.65E-10 | 1.2 | 0.3737 | 0.356 | NA | NA |
| *NOD2* | 16-50729867-G-GC | frameshift | rec | Union#K509#K50.9 Crohn's disease\| unspecified | 6.25E-10 | 31.5 | 0.0415 | 0.0179 | 0 | {Inflammatory bowel disease 1; Crohn disease} |
| *PALB2* | 16-23621362-C-T | stop gained | dom | 41202#BlockC50-C50#C50-C50 Malignant neoplasm of breast | 6.85E-10 | 6.9 | 0.0019 | 2.7E-04 | 1 | {Breast cancer; susceptibility to} |
| *IL33* | 9-6255967-G-C | splice acceptor | allele | 20002#1111#asthma | 1.02E-09 | 0.57 | 0.0026 | 0.0047 | NA | NA |
| *GCSAML* | 1-247556467-G-A | splice donor | allele | Union#L50#L50 Urticaria | 1.84E-09 | 1.3 | 0.0855 | 0.0678 | NA | NA |
| *BRCA2* | 13-32337160-TAAAC-T | frameshift | dom | 41202#Z40#Z40 Prophylactic surgery | 2.37E-09 | 565.1 | 0.0054 | 9.7E-06 | 1 | Multiple cancers |
| *APOBEC3B* | 22-38984097-C-T | stop gained | allele | Union#B029#B02.9 Zoster without complication | 2.90E-09 | 297.4 | 0.01 | 3.4E-05 | NA | NA |
| *CCDC198* | 14-57481662-T-C | splice acceptor | allele | Union#G43#G43 Migraine | 3.62E-09 | 0.90 | 0.2828 | 0.2625 | NA | NA |
| *CLECL1* | 12-9733111-A-ATAAGT | frameshift | allele | Union#E07#E07 Other disorders of thyroid | 8.79E-09 | 0.92 | 0.4748 | 0.4962 | NA | NA |

We investigated direction of effect consistency by selecting quantitative trait phenotypes with at least five individually significant rare non-synonymous variants (MAF<0.1%) in a given gene. For all the quantitative trait gene-phenotype relationships (17/17), we observed that ≥80% of the individually significant variants had the same direction of effect (**Fig 2e**). This observation indicates that it is uncommon to observe both negatively- and positively-associated rare variants for a specific gene-phenotype relationship (**Fig 2e**).

As expected, effect sizes observed for the collection of rare variants that were significantly associated with disease were substantially higher when compared to the collection of common variants (Wilcox *P* = 2.4 × 10^−40^) (**Fig. 2c**). While some of the ExWAS variants significantly associated with disease reflect linkage with nearby causal variants, the associated PTVs and missense variants themselves often, but not universally, represent functional candidates for such associations.^15^ Notably, 22% (65 out of 302) of the study-wide significant non-synonymous variants were either unreported in prior GWAS studies (including 50kbp flanking windows) or not labelled as Pathogenic / Likely Pathogenic in ClinVar (**Fig. 2d, Supplementary Table 2**). Collectively, these analyses of rare protein-coding variants introduce novel statistically significant genetic associations, many of which have higher effect sizes than previously established variants in existing catalogues.

### Rare variant collapsing analyses

In addition to variant-level association studies, we performed gene-level collapsing analyses. Rather than studying the effect of individual variants on disease, collapsing analysis assesses the effect of a class of variation on disease by comparing the aggregate number of cases to the aggregate number of controls that carry qualifying variants (QVs) in a given gene. This procedure produces one statistical test per gene instead of one per variant. Collapsing analysis has been shown to substantially increase the power to detect genetic risk in phenotypes driven by an allelic series.^17–20^ We performed collapsing analyses across 18,741 genes for 11,952 phenotypes and adopted 12 different QV classes (**methods and Supplementary Table 3**). This collection equated to 2.7 billion gene-phenotype tests. The 12 QV classes included 10 dominant models, one recessive model, and one synonymous variant model, akin to an empirical negative control. For each QV class, the proportion of cases is compared to the proportion of controls carrying QVs in each gene. The exception is the recessive model, in which a subject must carry two QVs or be hemizygous for a single QV (**methods**).

Due to high correlation among the 12 QV classes and among the assessed phenotypes, defining a significance threshold for this analysis poses a challenge. With a priority placed on avoiding false claims, we defined two null distributions: an empirical null distribution using the synonymous collapsing model, and an n-of-1 permutation based null distribution. These two null distributions independently converged on a study-wide significance threshold of *P* ≤ 5 ×10^−9^ (**methods**). Adopting this threshold as collapsing study-wide significance, we identified 424 significant gene-phenotype relationships for binary traits and 487 for quantitative traits (**Fig 3a, b, Supplementary Table 4**). The majority of these signals emerged from the PTV QV classes (76.4% of significant binary associations; 58.3% of significant quantitative associations), with the remaining signals attributable to classes expanding beyond PTVs, thus emphasising the importance of studying variant types beyond PTVs in a gene-based approach.

**Figure 3.**
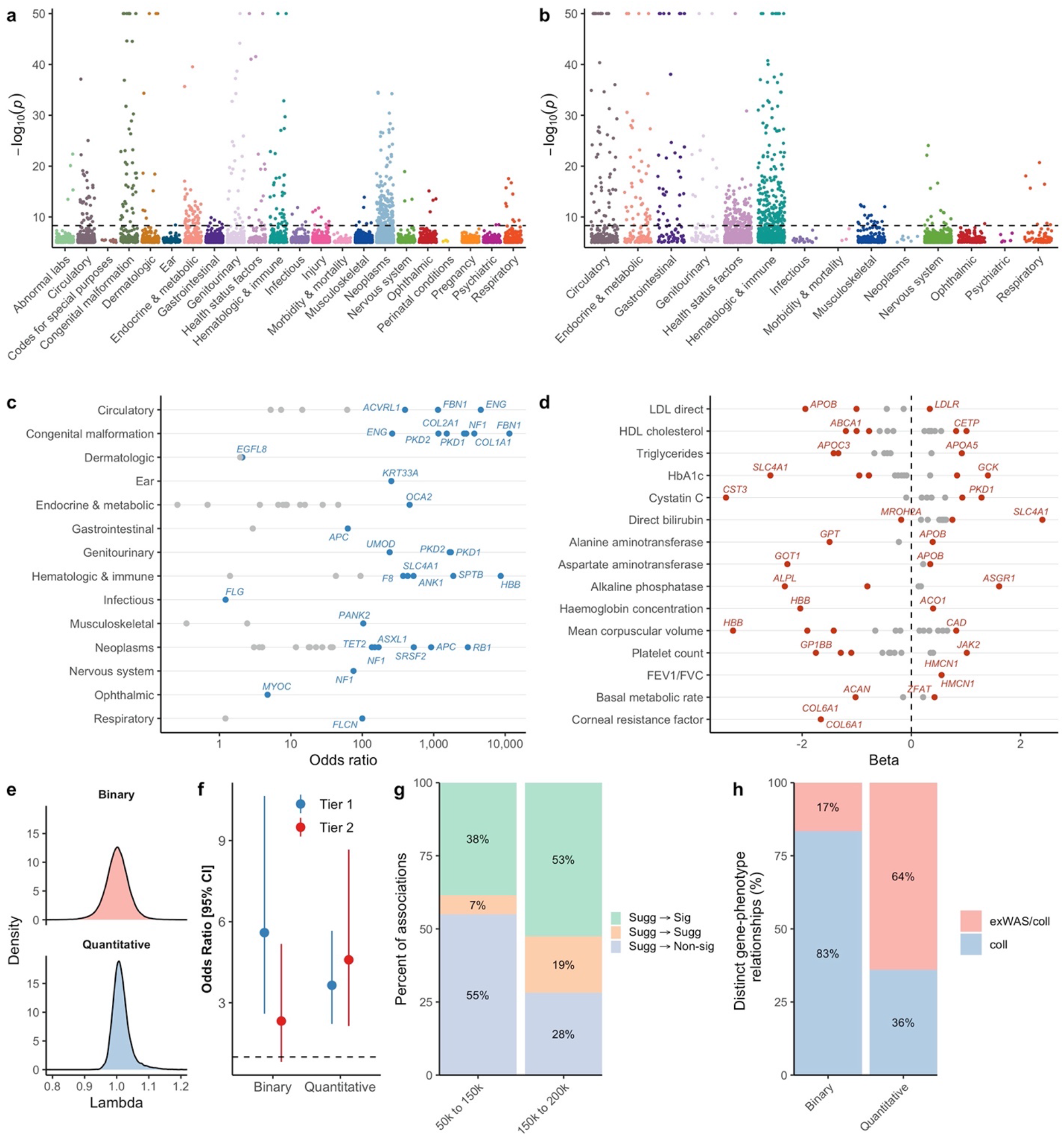
Summary of gene-level collapsing analysis results. **(a, b)** Manhattan plot depicting significant gene-phenotype associations for quantitative and binary traits, respectively. Only the strongest effect size association per collapsing model is displayed. The dashed line represents the genome-wide significant p-value threshold (5 × 10^−9^). The plots are capped at −log_10_p=50. **(c)** Depiction of select strong effect gene associations per disease area. Genes with the highest OR for a chapter or with OR>100 are labelled. **(d)** Illustration of large effect gene-phenotype associations for select disease-related biomarkers. **(e)** Distribution of lambda (inflation factor) values across all collapsing models for binary and quantitative traits**. (f)** Forest plot demonstrating enrichment for known drug targets stratified by statistical significance (Tier 1: statistically significant genes (*P* < 5×10^−^ 9); Tier 2: statistically suggestive genes (5×10^−9^ < *P* < 1×10^−7^)) achieved across all collapsing models for binary and quantitative traits. **(g)** Percentage of suggestive binary gene-phenotype associations that became significant (sig) (*P* < 5×10^−9^), non-significant (non-sig) (*P* > 1×10^−7^) or remained suggestive (sugg) (5×10^−9^ < *P* < 1×10^−7^) with each successive UKB tranche release for binary traits **(supplementary methods) (h)** Bar chart depicting the proportion of distinct gene-phenotype relationships detected either exclusively via collapsing analyses (coll) or through both single variant and collapsing analyses (ExWAS/coll).

The gene-level collapsing analyses identified a plethora of large effect signals across most disease areas and disease-relevant quantitative traits (**Fig. 3c, d**). The median genomic inflation factor (λ) across the collapsing analysis PheWAS was 1.002 for binary traits (range: 0.64-1.41) and 1.010 for quantitative traits (range: 0.92-1.33), indicating that our test statistics are highly robust to any systematic bias or other sources of inflation (**Fig. 3e**). The majority of the associations for binary phenotypes (84.9%) are supported by OMIM or as Pathogenic/Likely Pathogenic in ClinVar, providing further validation that our collapsing analysis paradigm captures robust rare variant-driven signals with high confidence (**Supplementary Table 4**). This includes well-established rare germline variants in genes associated with monogenic disease (for example, PTVs in polycystin 1 (*PKD1)* with chronic kidney disease and *HBB* with thalassemia), but also some genes with expanded phenotypes beyond those reported in OMIM. For example, if we look at the 12.0% of the European population who are carriers of a filaggrin (*FLG*) PTV, we find those carriers have significantly higher risk for well-known associations, such as dermatitis (*P*=3.9×10^−63^; OR: 1.95 [95%CI: 1.81 – 2.10]) and asthma (*P*=2.9×10^−^ 18; OR: 1.21 [95%CI: 1.16 – 1.27]),^21^ as well as additional diseases like melanoma (*P*=2.0×10^−7^; OR: 1.18 [95%CI: 1.11 – 1.26])^22^. Concomitant increases in vitamin D levels (*P*=4.8×10^−80^; β: 0.14 [95%CI: 0.13 – 0.16])^23^ suggest risk of melanoma and basal cell carcinoma in *FLG* PTV carriers could be attributable to increased sensitivity to ultraviolet B radiation. Such a resource presents the community with a powerful opportunity to investigate a wide spectrum of phenotypes associated with high-impact genetic aberrations in any given protein-coding gene of research interest.

Given the genotype-first approach employed across this large population-based dataset, penetrance estimates for gene-phenotype relationships can be generated with less ascertainment bias and greater confidence than those reported previously from case-based approaches. Penetrance estimates for rare (MAF<0.1%) PTVs in *FLG* with respect to asthma (15.5% [95% CI: 14.9% - 16.0%]) or eczematous dermatitis (4.5% [95% CI: 4.2% - 4.8%]) diagnoses contrast to penetrance estimates derived from gene-phenotype relationships of larger effects, such as PTVs in *HBB* and thalassaemia diagnoses (69.0% [95% CI: 49.2% - 84.7%]). It is foreseeable that through careful variant curation and more refined clinical phenotyping beyond simple ICD-10 codes, the UK Biobank dataset can potentially provide population-level penetrance estimates for established gene-disease relationships, not only for individual variants, but also for an aggregated class of variants (e.g., PTVs) in the gene. This might be limited to indications for which the prevalence in UK Biobank is consistent with that reported in literature.

Despite adopting a germline-tuned bioinformatics pipeline, seven study-wide significant genes were found to associate with haematological malignancies and this detection of somatic blood mutations was attributable to the blood-based sequencing nature of this dataset (**Supplementary Table 5 and Supplementary Table 6**). We also identified several genes that achieved study-wide significance for putatively protective PTVs, including classical examples of *APOB* and *PCSK9* (**Supplementary Table 7**) and more in the suggestive p-value range (**Supplementary Table 4**). Such signals provide validation for existing therapeutic strategies and may stimulate future therapeutic development opportunities.

Collectively, these rare-variant based findings reflect biological insight into common complex diseases and provide substrate for future therapeutic development opportunities. This is supported by the data that, for both binary and quantitative traits, UK Biobank derived statistically significant gene-phenotype associations were enriched for FDA approved drugs (binary OR: 5.60 [95% CI: 2.60-10.6]; *P*: 1.31×10^−6^; quantitative OR: 3.65 [95% CI: 2.22-5.66]; *P*: 5.18×10^−8^) (**Fig. 3f, methods**). Compared to two smaller ground-breaking UKB PheWAS’s that also applied gene-based aggregate statistics,^11,12^ we identified a 1.2 and 5.6-fold increase in unique statistically significant gene-trait associations using the same first tranche of 50K UKB data –– this can be attributed to the depth of outcomes studied and the differences in methodologies (**Supplementary Figure 1**).

It is also expected that many statistically suggestive (i.e., within 5 × 10^−9^ < *P* < 1 × 10^−7^ range) gene-phenotype associations represent true positive associations that remain relatively underpowered. We looked at the patterns across three UKB exome sequencing tranches and found that the proportion of suggestive associations for binary traits that achieve statistical significance in subsequent tranche is consistently high. For instance, approximately half of the suggestive associations reported in the 150K UKB exomes tranche achieved statistical significance in the 200K UKB exomes tranche (**Fig. 3g**).

We also found demonstrable complementarity between the variant-level ExWAS and gene-level collapsing analyses. Among the distinct gene-phenotype relationships identified through the collapsing analyses for the binary phenotypes, only 16.5% (70/424) were also detected using ExWAS, indicating that the remaining 83.5% were undetectable by traditional variant-level association tests applied on the same cohort (**Fig. 3h**). A noteworthy higher overlap was observed for quantitative phenotypes where 63.9% (311/487) of the collapsing analysis associations were detectable using variant-level ExWAS (**Supplementary Table 8**). This highlights that, particularly in the case-control scenario, a gene-based collapsing framework can identify associations that are currently undetectable by single variant-based approaches in an identical test setting, exemplifying the value of collapsing analyses for identifying additional biological insight and potentially novel drug targets. Focusing on the PTV models, we also examined the proportion of the statistically significant gene-trait associations identified using a more generous MAF filter of 5% (“ptv5pcnt” model) that were missed by both a stricter MAF threshold of 0.1% (“ptv” model) and the ExWAS analyses. We observe that 7.5% (21/281) and 10.2% (33/324) of gene-trait associations captured by the “ptv5pcnt” model were not captured by either the “ptv” model or ExWAS for quantitative and binary traits, respectively (**Supplementary Table 9**).

### Rare PTVs in *HMCN1* are associated with an increased FEV1/FVC ratio

Previously unreported PTV collapsing signals that emerge from quantitative physiological traits provide potential opportunities as novel therapeutic targets, but often require further functional investigation to understand their precise mechanism and disease relevance. One novel gene-phenotype signal emerging from the collapsing analysis was for the gene *HMCN1*, encoding Hemicentin 1 (**Fig. 4a**). Carriers of rare PTVs in *HMCN1* appear to have a significantly higher Forced Expiratory Volume in 1 second (FEV1) / Forced Vital Capacity (FVC) ratio z-score compared to non-carriers (**Fig. 4b**) (p = 2.0 × 10^−21^; beta = 0.55). Abnormalities in FEV1/FVC ratio may be seen in serious lung diseases, such as idiopathic pulmonary fibrosis (IPF), a fatal disorder characterised by progressive, destructive lung scarring, and rapid decline in FVC is associated with worsening survival.^24^ This disease process is thought to result from recurrent microinjury in alveolar epithelial cells that leads to aberrant repair and excess collagen and matrix production by myofibroblasts.^25^

**Figure 4.**
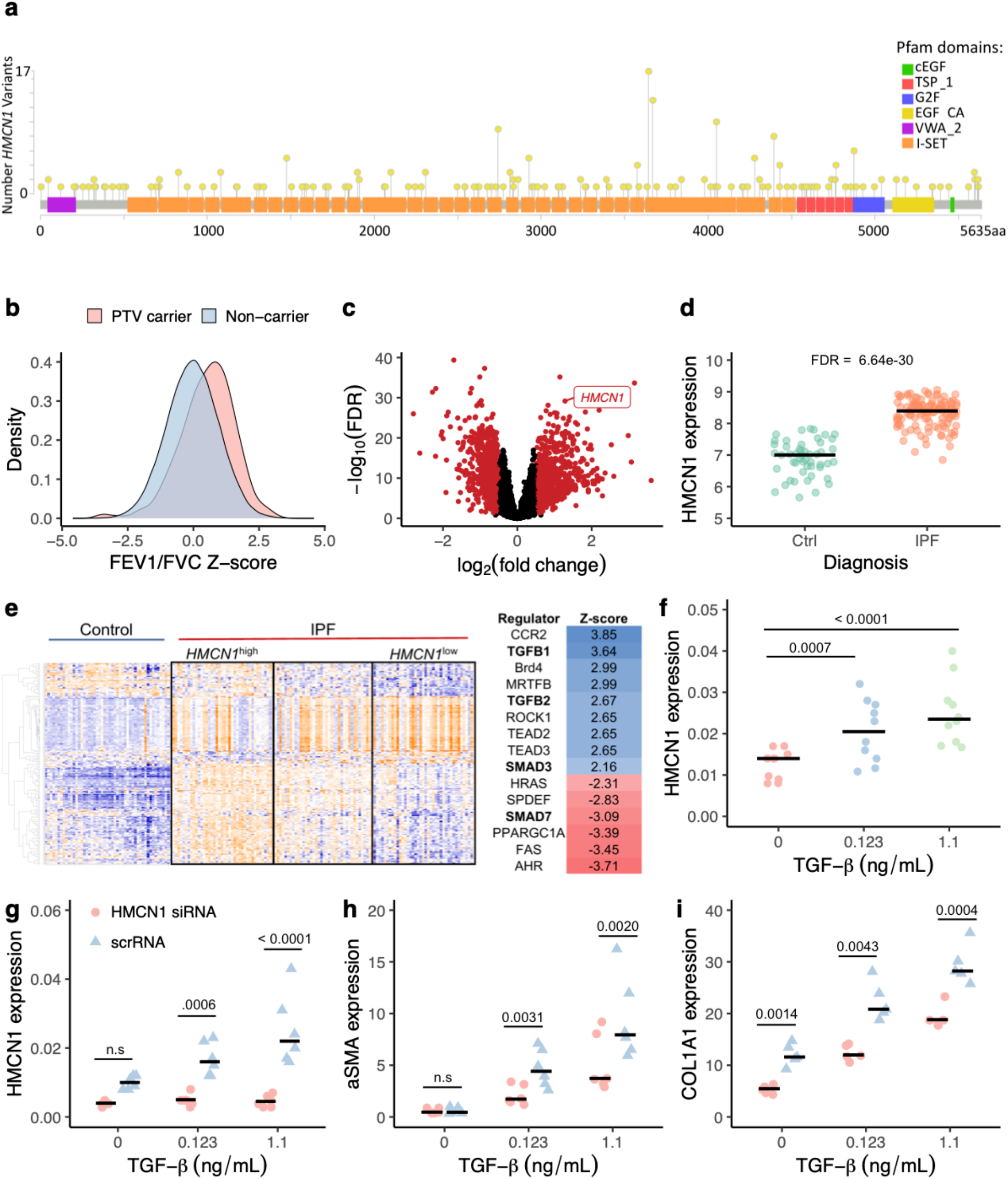
Functional evaluation of PTVs in *HMCN1*. **(a)** Locations of rare *HMCN1* PTVs among the UK Biobank participants. **(b)** The distribution of FEV1/FVC ratio Z-scores among *HMCN1* PTV carriers versus non-carriers. (**c**) Volcano plot depicting differentially expressed genes in IPF tissue versus control tissue. *HMCN1* is labelled. Red points indicate genes with an FDR < 0.05 and absolute log fold change > 0.5. **(d)** Expression of *HMCN1* in lung tissue derived from healthy controls versus patients with IPF. **(e)** Gene expression patterns of healthy lung tissue versus IPF tissue with high *HMCN1* expression and low *HMCN1* expression. Enriched up-stream regulators of genes differentially expressed between *HMCN1*^high^ and *HMCN1*^low^ IPF tissue are shown on the right. Bolded regulators are directly involved in TGFβ signaling. **(f)** HMCN1 expression in HFL1 (human foetal lung fibroblast) cells treated with TGFβ for 24h at different doses. **(g)** TGFβ-induced expression of HMCN1 in HFL1 cells treated with either *HMCN1* siRNA or scrambled RNA control (scrRNA). **(h)** Alpha-smooth muscle actin (aSMA) expression in cells treated with *HMCN1* siRNA versus scrRNA. Expression of COL1A1 in cells treated with siRNA versus scrRNA.

To better understand the biological mechanism underlying the *HMCN1*-lung function association, we leveraged RNA microarray data derived from 167 individuals with IPF and 50 non-diseased controls (GSE32537).^26^ We found that *HMCN1* expression was significantly increased in tissue derived from IPF patients and ranked among the top 150 out of 835 significantly upregulated genes. This is consistent with the human genetic data (**Fig. 4 c,d**). Patients with the highest *HMCN1* expression exhibited a distinct gene expression pattern when compared to patients with low *HMCN1* expression (**Fig. 4e**). We performed a pathway level analysis to predict regulators of the genes differentially expressed between these two groups. Among the top 15 predicted up-stream regulators, four were directly related to TGFβ signalling: *TGFB1*, *TGFB2, SMAD3,* and *SMAD7* (**Fig. 4e**). *TGFB1* had an enrichment z-score of 3.64 (p=3.3×10^−12^), suggesting that differential expression patterns in individuals with high *HMCN1* expression may be driven by *TGFβ* signalling (**Fig. 4e**).

To test for a direct association between TGFβ signalling and HMCN1 expression, we treated human foetal lung fibroblasts with TGFβ and found that HMCN1 expression increased in a dose-dependent manner (**Fig. 4f**). Previous evidence indicates that TGFβ plays an important role in differentiating fibroblasts into myofibroblasts by stimulating the expression of alpha-smooth muscle actin (αSMA). We found that upon siRNA knockdown of *HMCN1* (**Fig. 4g**), treatment with TGFβ1 resulted in significantly reduced αSMA expression compared to controls (**Fig. 4h**). *HMCN1* knockdown also resulted in reduced expression of collagen 1 (COL1A1), an important component of extracellular matrix deposition seen in IPF (**Fig. 4i**). Thus, *HMCN1* may be required for this profibrotic process regulated via the TGFβ signalling axis.

Collectively, our results begin to unravel how the presence of PTVs in *HMCN1* may affect lung function and that the increased expression of this gene may be involved in IPF pathogenesis. PTVs in *HMCN1* are not significantly associated with other disease-related traits among our UKB PheWAS, suggesting that inhibiting this gene could be a tolerable therapeutic intervention for IPF and potentially other disorders of the lung.

## Discussion

Using exome sequencing of 177,882 unrelated European ancestry UKB participants combined with records of 11,952 phenotypes, we have performed a pheWAS of unprecedented scale and identified 31,184 variant- and 911 gene-level statistically significant phenotypic relationships. The latter offers the advantage of providing a clearer link between causal gene(s) and phenotype(s). Our variant-level association tests include variants that are not frequent enough to be captured by previous microarray-based genotyping studies. We also applied gene-level collapsing analyses to test the aggregate effect of private-to-rare functional variants for which single-variant approaches are inadequate. The majority of significant gene-level associations were not captured by variant-level analysis, demonstrating the complementarity of the two approaches. Many of the study-wide significant associations have been previously reported, supporting the robustness of our results. Furthermore, in all instances where a particular gene harboured multiple rare variants significantly associated with a quantitative trait phenotype, majority (≥80%) of those variants had the same direction of effect.

Using 177,882 UKB samples has increased (12-fold) our ability to detect study-wide significant associations compared to when adopting the original 50K UKB exomes. The scale of the study also presents challenges like defining an appropriate significance threshold. Here we show how this could be addressed using an empirical null distribution as a negative control model alongside an n-of-1 permutation.

We found that the study-wide significant gene-phenotype associations are significantly enriched for targets of FDA-approved drugs. Additionally, among our novel findings is the association of PTVs in *HMCN1* with lung function, suggesting that *HMCN1* could become a potentially promising candidate drug target for respiratory disease. These two findings highlight the utility of mining UK Biobank PheWAS outputs as a source of high impact genetic support for target identification and evaluation, which when followed up with functional investigation to understand the underlying biological and disease mechanisms, can help improve the efficiency of pharmaceutical pipelines.^5,27^

As a consequence of the recruitment criteria for the UKB, the richest yields of this study relate to common human diseases and routinely measured quantitative traits such as biomarkers rather than severe early-onset diseases. Beyond the PheWAS setting, future refinement of phenotypic definitions by combining binary and/or quantitative phenotypic information, alongside any temporal data, may better reflect disease heterogeneity. Moreover, this study is currently limited to SNVs and indels, although the utility of evaluating copy number variants has been demonstrated by others.^28^ While our dataset is limited to individuals of European ancestry, we also highlight the need to use UK Biobank as a gold standard and establish equivalent resources for other global populations. Furthermore, the exogamous nature of this cohort meant that we detected homozygous PTVs—which can provide an *in vivo* model to study the phenotypic effect of gene knockout^29^—for only ~18% genes.

Altogether, this PheWAS evaluating the genetic architecture of human disease yields a rich resource of statistically robust high-impact gene-phenotype associations at an unprecedented scale, with the potential to elucidate novel disease mechanisms, identify phenotypic expansions for known disease genes and enable the continued development of novel human genetics-validated drug targets.

## Methods

### 1. UK Biobank Resource

The UK Biobank is a prospective study of approximately 500,000 participants aged 40-69 years at recruitment. Participants were recruited in the UK between 2006-2010 and continuously followed.^30^ Participant data includes health records that are periodically updated by the UK Biobank, self-report survey information, linkage to death and cancer registries, collection or urine and blood biomarkers, imaging data, accelerometer data and various other phenotypic endpoints.^7^

### 2. Phenotypes

We studied two main phenotypic categories: binary and continuous traits taken from the February 2020 data release that was accessed on March 27^th^ 2020 as part of UKB application 26041. We then updated Hospital Episode Statistic (HES) and death registry data using the *ad hoc* release by the UK Biobank on July 2020.

To parse the UKB phenotypic data we adopted a modified version of the PHESANT package that can be located at https://github.com/astrazeneca-cgr-publications/PEACOK. The adopted parameters are available in **Supplementary Methods** and have been previously introduced in PHESANT (https://github.com/MRCIEU/PHESANT).^13^

For UK Biobank tree fields, such as the ICD10 hospital admissions (Field 41202), we studied each leaf individually, and also studied as separate phenotypic entities each of the groupings up to the highest level of the ICD10 root chapter phenotypes.

Furthermore, for the tree-related fields (Fields: 20001, 20002, 40001, 40002, 40006 and 41202), to reduce the potential contamination of potentially genetically-related diagnoses among controls we restricted controls to subjects who did not have a positive diagnoses in that corresponding chapter. A minimum of 20 cases were required for a binary trait to be studied.

In addition to studying UK Biobank algorithmically-defined outcomes, we also constructed a union phenotype for each ICD10 phenotype. These union phenotypes are denoted by a “Union” prefix and the applied mappings are available in **Supplementary Table 1**.

In total, we studied 10,533 binary and 1,419 continuous phenotypes. For all binary phenotypes we gender matched controls when the % female cases was significantly different (Fisher’s exact p < 0.05) to the % available female controls. This included sex-specific traits where, by design, all controls would be same sex as cases. Finally, to allow for more compartmentalised ICD10 chapter-based analyses, all 11,952 binary and quantitative trait phenotypes were mapped to a single ICD10 chapter including manual mapping for the non ICD10 phenotypes. These chapter mappings are provided in **Supplementary Table 1**. It is acknowledged that chapter mapping may have greatest utility for diagnostic, rather than procedural, ICD10 codes. For procedural codes, genetic associations could be incorrectly interpreted if chapter mappings are relied on. For example, surgical procedures commonly performed for oncology patients are categorized within the dermatology chapter. Genetic associations reported for these procedures would be categorized within the dermatology chapter, but the underlying disease process is instead most likely reflective of an oncology aetiology.

### 3. Sequencing

Whole-exome sequencing (WES) data for UK Biobank participants were generated at the Regeneron Genetics Center (RGC) as part of a pre-competitive data generation collaboration between AbbVie, Alnylam Pharmaceuticals, AstraZeneca, Biogen, Bristol-Myers Squibb, Pfizer, Regeneron and Takeda with the UK Biobank.^31^ Genomic DNA underwent paired-end 75bp whole exome sequencing (WES) at Regeneron Pharmaceuticals using the IDT xGen v1 capture kit on NovaSeq6000 machines. Exome sequences from 200,593 UK Biobank participants were made available to the Exome Sequencing consortium. Initial QC was performed by Regeneron and included sex discordance, contamination, unresolved duplicate sequences and discordance with microarray genotyping data checks, as previously described.^12^

### 4. AstraZeneca Centre for Genomics Research (CGR) Bioinformatics Pipeline

The 200,593 UKB exome sequences were processed at AstraZeneca from their unaligned FASTQ state. We adopted an in-house cloud compute platform running Illumina DRAGEN Bio-IT Platform Germline Pipeline v3.0.7 to align the reads to the GRCh38 genome reference and perform small variant SNV and indel calling. SNVs and indels were annotated using SnpEFF v4.3 against Ensembl Build 38.92. We further annotated all variants with their gnomAD minor allele frequencies (gnomAD v2.1.1 mapped to GRCh38).^10^

An additional two bioinformatic scores were adopted specifically for missense variants. We adopted missense tolerance ratio (MTR) scores to permit focused analyses of missense variants occurring in the most missense constraint regions of individual human protein-coding genes, as calculated based on the gnomAD reference cohort.^32^ We also adopted the REVEL score as bioinformatic tool to support further prioritisation of deleterious predicted missense variants.^33^

### 5. Additional Quality Control

In addition to what had already been QC flagged, we excluded from our analyses 77 (0.04%) sequences that achieved a VerifyBAMID freemix (contamination) level >4%,^34^ and an additional five sequences (0.002%) where <94.5% of the consensus coding sequence (CCDS release 22) achieved a minimum of 10-fold read-depth.^35^

To mitigate against possible bias driven by relatedness, we performed pair-wise relatedness checks across all remaining UK Biobank exome sequenced participants. This was achieved by estimating pairwise kinship coefficients using the --kinship algorithm of KING v2.2.2^36^ and exome sequence-derived genotypes for 43,889 biallelic autosomal SNVs located in coding regions.

The ukb_gen_samples_to_remove() function from the R package ukbtools^37^ was used to choose a subset of individuals within which no pair had a kinship coefficient exceeding 0.0884. This function aims to choose a maximal set using a greedy algorithm that iteratively selects for removal among pairs of relatives above this threshold the member of each pair with the highest number of relatives above the threshold. Through this process, an additional 9,489 (4.73%) sequences were removed from downstream analyses.

This process resulted in a remaining collection of 191,022 (95.2%) UK Biobank unrelated sequences of any genetic ancestry that were available for analyses presented in this work.

### 6. Genetic Ancestry

For case-control cohort analyses we further restricted the statistical tests to reflect a homogeneous European genetic ancestry test cohort. This was achieved by running the exomes through PEDDY v0.4.2 with the ancestry labelled 1K Genomes Project reference sequence data for genetic ancestry predictions. Of the above 191,022 UK Biobank unrelated sequences, 12,784 (6.4%) achieved a Pr(European) ancestry prediction <0.99. Focusing on the remaining 178,238 UK participants we further restricted the European cohort to those within ±4 standard deviations across the top four principal component means resulting in an additional 356 (0.2%) outlier participants. The remaining collection of 177,882 (88.7%) unrelated European ancestry UK Biobank participants were adopted for all case-control analyses reported in this manuscript.

### 7. ExWAS Analyses

We tested variants observed among at least six individuals from the 177,882 unrelated European UK Biobank exomes. Variants were required to pass the following QC criteria: minimum coverage 10X; percent of alternate reads in heterozygous variants ≥ 0.2; binomial test of alternate allele proportion departure from 50% in heterozygous state p > 1×10^−6^; genotype quality score (GQ) ≥ 20; Fisher’s strand bias score (FS) ≤ 200 (indels) ≤ 60 (SNVs); mapping quality score (MQ) ≥ 40; quality score (QUAL) ≥ 30; read position rank sum score (RPRS) ≥ −2; mapping quality rank sum score (MQRS) ≥ −8; DRAGEN variant status = PASS; variant site is not frequently missing (i.e., <10X coverage) in ≥ 10% of sequences; variant did not fail any of the aforementioned QC in ≥ 5% of sequences; variant site achieved 10-fold coverage in ≥ 30% of gnomAD exomes, and if variant was observed in gnomAD exomes, ≥50% of the time those variant calls passed the gnomAD QC filters (gnomAD exome AC / AC_raw ≥ 50%).

To account for large case-control imbalances and permit the robust study of extremely rare occurring variants (as low as 6 allele observations, i.e., MAF > 0.0017%), variant-level p-values were generated adopting a Fisher’s exact test. Three distinct genetic models were studied for the binary traits: allelic (A vs B allele), dominant (AA+AB vs BB) and recessive (AA vs AB+BB), where A denotes the alternative and B denotes the reference allele. For the continuous traits, the allelic model is replaced with a genotypic (AA vs AB vs BB) test. For ExWAS analysis we adopted the significance cutoff of p<1×10^−8^. ^38^

### 8. Collapsing Analyses

To perform collapsing analyses, we aggregate variants within each gene that fit a given set of criteria, identified as “qualifying variants” (QVs).^20^ Overall, we performed 11 non-synonymous collapsing analyses, including 10 dominant and one recessive model, plus an additional synonymous variant model as an empirical negative control. In each model, for each gene, the proportion of cases is compared to the proportion of controls carrying one or more qualifying variants in that gene. The exception is the recessive model, where a subject must have two qualifying alleles, either in homozygous or potential compound heterozygous form. Hemizygous genotypes for the X chromosome were also qualified for the recessive model. The criteria behind the QV classifications for each collapsing analysis model are in **Supplementary Table 3**. To account for large case-control imbalances and permit the robust study of sparse observations, collapsing analysis p-values were generated adopting a Fisher’s exact test.

For all models (**Supplementary Table 3**) we applied the following QC filters: minimum coverage 10X; annotation in CCDS transcripts (release 22; ~34Mb); percent of alternate reads in heterozygous variants ≥ 0.25 and ≤ 0.8; binomial test of alternate allele proportion departure from 50% in heterozygous state p > 1×10^−6^; genotype quality score (GQ) ≥ 20; Fisher’s strand bias score (FS) ≤ 200 (indels) ≤ 60 (SNVs); mapping quality score (MQ) ≥ 40; quality score (QUAL) ≥ 30; read position rank sum score (RPRS) ≥ −2; mapping quality rank sum score (MQRS) ≥ −8; DRAGEN variant status = PASS; variant site achieved 10-fold coverage in ≥ 25% of gnomAD exomes, and if variant was observed in gnomAD exomes the variant achieved exome z-score ≥ −2.0 and exome MQ ≥ 30.

### 9. Defining the study-wide significant cut-offs for collapsing analyses

Given the high degree of correlation among the studied phenotypes and also the high degree of similarity among the multiple collapsing models applied there was a need to define a more appropriate study-wide significance threshold for gene-level PheWAS as Bonferroni correction is inappropriate in this PheWAS context. We took two approaches to define the study-wide significance thresholds for the gene-based collapsing PheWAS.

One of the collapsing analysis models that was applied as an empirical negative control was the synonymous model. Here, it is expected that in general, synonymous variants will not have a significant contribution to disease risk and could thus act as a useful empirical negative control for study-wide p-value thresholding. Surveying across the 10,533 studied binary phenotypes and considering the 18,741 studied genes we had a distribution of 197,398,953 Fisher’s Exact test statistics corresponding to the synonymous collapsing model. Among the tail of this distribution for binary traits we observe a tail of p-values beginning from p=3.8×10^−9^ (Supplementary Table 10). Similarly for the 1,419 quantitative phenotypes we had a distribution of 26,593,479 Fisher’s Exact test statistics corresponding to the synonymous collapsing model. Among the tail of this distribution we identify a single genuine relationships: *MACROD1* synonymous variants correlating with decreased levels of ‘Urate’ (p=8.6×10^−15^).^39^ Following this known relationship, we see a trail of p-values beginning from p=1.3×10^−7^ (Supplementary Table 10).

Given this magnitude of test statistics generated in PheWAS scale another proposal has been the utility of a n-of-1 permutation.^40^ Here, we shuffled the case-control (or quantitative measurement) labels once for every phenotype while maintaining the participant-genotype structure and performed an n-of-1 permutation across all 11 non-synonymous collapsing models for the binary traits (2,171,388,483 tests) and the quantitative traits (292,528,269 tests). Reviewing the tails of these two n-of-1 permutation-based p-value distributions the lowest permutation-based p-value achieved was 4.2×10^−9^ (binary tests) and 9.6×10^−9^ (quantitative tests).

Given the scale and correlations among this dataset, we found both these approaches provide suitable alternatives to the Bonferroni p-value threshold, which in this case would be p<2.0×10^−11^. Guided by both empirical and n-of-1 permutation based null p-value distributions we define a study-wide significance cut-off of p≤5×10^−9^ for the non-synonymous collapsing analysis results presented in this manuscript (Supplementary Table 10).

Finally, for each of the 143,424 exome-wide collapsing analyses comprising the collapsing PheWAS (12 models * [10,533 + 1,419] studied phenotypes) we calculated the lambda genomic inflation factor (λ) after excluding genes achieving exome-wide significance p<2.6×10^−6^ for that phenotype.

### 10. Collapsing analysis enrichment for approved drug targets

Drug target analysis was performed using a publicly available list of genes (n=387) considered to represent drug targets (source: https://raw.githubusercontent.com/ericminikel/drug_target_lof/master/data/drugbank/drug_gene_match.tsv), that were originally derived from DrugBank.^41^ For each gene, the most statistically significant collapsing analysis phenotypes were identified, before being partitioned into three categories (significant (*P* < 5×10^−9^) (binary n=87, quantitative n=291), suggestive (5×10^−9^ < *P* < 1×10^−7^) (binary n=108, quantitative n=104), or non-significant (*P* > 1×10^−7^) (binary n=18,551, quantitative n=18,351)). The relationship between drug target status and gene-phenotype significance was assessed using a logistic regression model in R v3.5.1.

## ACKNOWLEDGEMENTS

We thank the participants and investigators in the UK biobank study who made this work possible (Resource Application Number 26041). We thank the UK Biobank Exome Sequencing Consortium (UKB-ESC) members AbbVie, Alnylam Pharmaceuticals, AstraZeneca, Biogen, Bristol-Myers Squibb, Pfizer, Regeneron and Takeda for funding the generation of the data and Regeneron Genetics Center for completing the sequencing and initial quality control of the exome sequencing data. We thank the AstraZeneca Centre for Genomics Research Analytics and Informatics team for processing and analysis of sequencing data. We thank Dr Matthew Hurles for valuable feedback on this manuscript.

