## Supplemental text for "Surveying the contribution of rare variants to the genetic architecture of human disease through exome sequencing of 177,882 UK Biobank participants"

### Supplementary Methods: Parameters adopted from PHESANT package

make_option(c("--catmultcutoff"), type="integer", default=10,
help="The cutoff for exclusion when creating dichotomous variables for CAT-multiple."),

make_option(c("--catordnacutoff"), type="integer", default=500,
help="The cutoff for exclusion for number of non-NAs in ordered categorical variables."),

make_option(c("--catunordnacutoff"), type="integer", default=500,
help="The cutoff for exclusion for number of non-NAs in unordered categorical variables."),

make_option(c("--contnacutoff"), type="integer", default=500,
help="The cutoff for exclusion for number of non-NAs in continuous variables."),

make_option(c("--binnacutoff"), type="integer", default=500,
help="The cutoff for exclusion for number of non-NAs in binary-variables."),

make_option(c("--bintruecutoff"), type="integer", default=30,
help="The cutoff for exclusion for numbers of members of a category in binary-variables."),

make_option(c("--mincategorysize"), type="integer", default=10,
help="The minimum number of samples in a category for categorical single, integer, and continuous variables."),

make_option(c("--maxunorderedcategories"), type="integer", default=1000,
help="The maximum number of categories in an unordered categorical variable"),

make_option(c("--propforcontinuous"), type="double", default=0.2,
help="The cutoff for proportion of samples with the same value for the variable to not be considered continuous.")

### Supplementary Methods: Informativeness of individual genes

Observing and equally importantly not observing signals among both gene- and variant-level association statistics are highly dependent on how informative the sequence data is for those regions of the exome.

We identified that the exome sequence data for UK Biobank participants who contributed to the analyses represented in this paper had on average 97.1% of the 34.1Mbp of the consensus coding sequence (CCDS release 22) covered with at least 10-fold coverage. To determine how informative individual genes were, we further broke down this assessment to the 18,741 genes represented in CCDS release 22. Overall, we found that 2,703 (14.4%) of genes had on average 100% of the protein-coding sites covered with at least 10x coverage among all UKB participants. A further 14,831 (79.1%) of genes had on average ≥99% of the protein-coding sites covered with at least 10x coverage among the UKB participants. An additional 884 (4.7%) of genes had on average ≥75% of the protein-coding sites covered with at least 10x coverage among the UKB participants. Leaving a remainder of 323 (1.7%) CCDS release 22 genes with < 75% of the protein-coding sites covered with at least 10x coverage among UKB participants. The individual gene-level coverage statistics are available in **Supplementary Table 11** and are a valuable resource for understanding the potential blind-spots in the human exome among this collection.

### Supplementary Methods: Evaluating suggestive signals in subsequent tranches

Exome sequence data was obtained in three discrete batches, with the cumulative total number of available exomes increasing from 50k to 150k, and subsequently 200k. A *P* threshold of *<* 5x10^-9^ was considered significant, suggestive was defined as 1x10^-7^ < *P* < 5x10^-9^ and non-significant reported as *P* > 1x10^-7^. All model-specific gene-phenotype relationships were harmonised across three tranches (50k, 150k, and 200k). The outcome of gene-phenotype-model relationships that appeared suggestive (i.e. 1x10^-7^ < *P* < 5x10^-9^) in the *i^th^* tranche were evaluated in the *i+1* tranche.

### Supplementary Methods: Comparing gene-level results of 50K UKB across multiple published studies

The first tranche of 50,000 UKB exomes was initially analysed by Van Hout et al under a collaborative effort between Regeneron Genetics Centre and GlaxoSmithKline,^1^ and later on by Cirulli et al within the auspices of Helix (https://ukb.research.helix.com/).^2^ Both groups performed gene-level collapsing analyses to explore the aggregate effect of rare variants on thousands of binary and quantitative traits found in the UKB repository. Their and our efforts differ in terms of the choice of variants collapsed, filters such as MAF, choice of phenotypes studied and statistical models. Here we endeavour to list the differences in methodology and subsequent results from the three efforts.

Van Hout et al^1^ used 49,960 individuals of European ancestry and focused on autosomal genes with > 3 predicted loss of function (pLoF) variants with MAF ≤ 1%. They run an additive collapsing model such that any individual that is heterozygous for at least one qualifying LoF in that gene region is considered heterozygous, and individuals that carry two copies of the same LoF are considered homozygous. Quantitative measures with ≥ 5 individuals were rank-based inverse normal transformed and analysed using BOLT-LMM v2.^3^ Prior to normalization, traits were first transformed as appropriate (log10, square) and adjusted for a standard set of covariates including age, sex, study site, first four principal components of ancestry, and in some cases BMI and/or smoking status. Data-points greater than five median absolute deviations from the median were excluded as outliers prior to normalization. Binary traits were based on ICD-10 diagnosis (primary diagnosis or ≥ 2 secondary diagnoses in in-patient Health Episode Statistics records), self-reported illness from verbal interview and physician-diagnosed illness. Binary outcomes with ≥ 50 cases were assessed with covariate adjustment for age, sex and first four principle components of ancestry using a generalized mixed model implemented in SAIGE.^4^

Cirulli et al^2^ used a dominant model and collapsed: 1) LoF variants only, and 2) coding variants only and not Polyphen or SIFT benign. The qualifying variants had MAF < 0.1% in the European ancestry UK Biobank unrelated set, as well as in any gnomAD population. Phenotypes were processed using the Neale lab modified version of PHESANT^5^. Statistical analysis was performed by BOLT-LMM,^3^ adjusting for age and sex. Genes needed to have at least five carriers of qualifying variants for quantitative traits and at least ten carriers of qualifying variants to be expected in the smaller sample group for binary traits. The traits that failed to run with BOLT-LMM (trait heritability fell below the required algorithm threshold) were analysed by linear regression using plink (quantitative traits adjusted for age, sex and the first 10 European-specific principle components) and Fisher’s exact test (binary traits) in the subset of unrelated European ancestry individuals. The Fisher’s exact test was also used to identify associations when <10 cases were expected to be carrying qualifying variants based on the overall prevalence. Although Cirulli et al^2^ analysed all ancestries on top of the European ancestry-only subset, here we focused on the results using European ancestry only.^2^

Focusing on the initial 50K exomes compared to these two ground-breaking UKB PheWAS studies^1,2^ on the initial 50K exomes, we found 62 statistically significant (*P* < 3.4x10^-10^ adopted by ^2^) gene-trait pairs (5 for binary and 57 for quantitative outcomes), while Van Hout et al^1^ identified 11 gene-trait pairs (2 binary and 9 quantitative) and Cirulli et al^2^ reported 51 gene-trait pairs (7 binary and 44 quantitative phenotypes) (Tables 5 & 6 from ^1^; Supplementary Data 2 from ^2^; **Supplementary Table 12**). Across the 81 unique statistically significant gene-trait associations found among phenotypes analysed by the three studies, 24 were exclusively identified in this study, 14 exclusively by Cirulli et al^2^ and 5 exclusively by Van Hout et al^1^ (**Supplementary Figure 1; Supplementary Table 12**).

### Supplementary Methods: *HMCN1* functional experiments

#### Expression analysis

We reprocessed array data from GEO dataset GSE32537, which includes 167 subjects with idiopathic pulmonary fibrosis and 50 non-diseased controls.^6^ The CEL files were processed using the robust multichip average (RMA) method^7^ along with BrainArray (<http://brainarray.mbni.med.umich.edu/Brainarray/Database/CustomCDF/genomic_curated_CDF.asp>) ensembl gene ID version 24 to collapse the probes to a single gene. This generated normalised expression data on a log_2_ scale with probes collapsed to gene level. QC indicated one sample did not correspond to the assigned gender in the metadata and was removed from subsequent tests.

**Differential gene expression:**

Differentially expressed genes were generated by using Limma moderated t-test in Array Studio to compare the IPF samples to the controls controlling for both age and gender as the controls were generally younger than IPF patients. We considered genes with an FDR < 0.05 and absolute log_2_ fold change > 0.5 as significantly differentially expressed.

In a separate analysis, we performed differential gene expression between IPF samples who had high versus low expression of *HMCN1.* IPF samples were divided into terciles based on their *HMCN1* expression levels (high, med, low). Differentially expressed genes for the were generated using Limma moderated t-test^8^ in Array Studio to compare the HMCN1^high^ samples to the HMCN1^low^ samples. Hierarchical clustering (Pearson correlation distance and complete linkage) was performed using the differentially expressed genes to generate a heatmap of all samples grouped according to HMCN1 expression group.

Functional enrichment was performed on the differentially expressed genes using IPA (QIAGEN Inc., https://www.qiagenbioinformatics.com/products/ingenuitypathway-analysis) using the Core Analysis tool. The Disease and Functions results with p-value <0.05 were extracted for further filtering. The activation status of the functions/pathways were predicted using IPA Upstream Regulator Analysis Tool by calculating a regulation Z-score and an overlap p-value, which were based on the number of known target genes of interest pathway/function, expression changes of these target genes and their agreement with literature findings. The detailed descriptions of IPA analysis are available under “Downstream Effects Analysis” on the IPA website (https://www.qiagenbioinformatics.com/products/ingenuitypathway-analysis).

#### Cell culture

Human foetal lung fibroblasts (HFL1) cells were cultured in MEM/ HEPES/ Glutamax^TM^ medium (ThermoFisher Scientific) supplemented with 10% FCS, penicillin and streptomycin. For TGF stimulation cells were seeded in 48 well plates at 4,5x10^4^ cells/well in 500ul and cultured for 24h followed by 24h culture in serum free medium (MEM/ HEPES/ GlutaMAX^TM^ medium supplemented with penicillin and streptomycin). Finally, cells were treated with TGF at a final concentration of 0.123ng/ml or 1,1 ng/ml or left untreated as control. 24 hours after stimulation cells were harvested for analysis.

#### siRNA transfection

HMCN1 knock down in HFL1 cells was achieved performing reverse transfection using Silencer® Select siRNA (final concentration 10nM, ThermoFisher Scientific s38249) and Lipofectamine RNAiMAX transfection reagent (1.5ul/ well, ThermoFisher Scientific). Silencer® Select negative control (ThermoFisher Scientific 4390843) was used as control siRNA. RNAiMax and siRNA were diluted in Opti-MEM reduced serum medium with GlutaMAX^TM^ (ThermoFisher Scientific) in 48 well plates before seeding of cells as described above. 24 hours after transfection cells were starved for another 24 hours followed by 24 hour TGF treatment.

#### RNA isolation and Real Time PCR

After TGFb treatment cells were washed twice in PBS before lysis in 100ul RLT buffer (Qiagen). RNA isolation was performed using RNeasy 96 well kit (Qiagen) according to the manufactures instructions. High-Capacity cDNA reverse transcription kit (ThermoFisher Scientific) was used for cDNA synthesis followed by Real Time PCR using TaqMan Gene Expression Assays (HMCN1 Hs00913222_m1, Col1A1 Hs00164004_m1, ACTB Hs01060665_g1, HPRT Hs02800695_m1 ThermoFisher Scientific).

#### Statistical analysis

P-values were generated using repeated measurement one-way ANOVA in Prism GraphPad v8.4.2.

Supplementary Figure 1: Venn diagram for gene-trait associations identified by three studies using the first tranche of 50K UKB.

There are 81 unique significant gene-trait associations (*P* < 3.4x10^-10^) found among phenotypes studied across the three efforts.

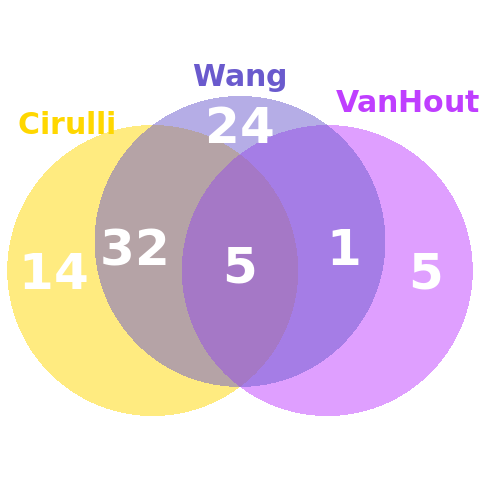

### Supplementary Table 1 – Studied phenotypes

Provided as separate additional file

Details of all phenotypes studied in this dataset. Binary: 10,533 binary phenotypes used for both gene-level collapsing analysis and variant-level ExWAS. Path and Root refer to UKB data structure as described in Showcase here <https://biobank.ndph.ox.ac.uk/showcase/browse.cgi>. Quantitative: 1,419 continuous traits used for both gene-level collapsing analysis and variant-level ExWAS.

### Supplementary Table 2 – ExWAS top hits

Provided as separate additional file

Significant genotype-phenotype associations from variant-level exome-wide association study. ExWAS_binary: binary associations (all variant consequences) excluding MHC region (chr6:25Mbp-35Mbp) with p<1x10^-8^. ExWAS_binary_non-syn: The subset of ExWAS_binary that are non-synonymous variants with p<1 x10^-8^. ExWAS_quantitative: quantitative associations (all variant consequences) excluding MHC region (chr6:25Mbp-35Mbp) with p<1 x10^-8^.

### Supplementary Table 3 – Collapsing analysis models

| Collapsing model | GnomAD MAF* | UKB MAF | UKB cohort no call or QC fail^ | Variant type | REVEL^11^ cutoff | Missense Tolerance Ratio (MTR)^12^ cutoffs |
| --- | --- | --- | --- | --- | --- | --- |
| **syn** (synonymous negative control) | ≤ 0.005% | ≤ 0.05% | ≤ 0.005% | Synonymous | - | - |
| **ptv** (Protein Truncating) | ≤ 0.1% (popmax) | ≤ 0.1% | ≤ 0.01% | PTV | - | - |
| **ptv5pcnt** (Protein Truncating, ≤5% MAF) | ≤ 5% (popmax) | ≤ 5% | ≤ 0.5% | PTV | - | - |
| **UR** (Ultra-rare damaging) | 0% | ≤ 0.005% | ≤ 0.001% | Non-synonymous | ≥ 0.25 | - |
| **URmtr** (Ultra-rare damaging, MTR informed) | 0% | ≤ 0.005% | ≤ 0.001% | Non-synonymous | ≥ 0.25 | MTR ≤ 25^th^ %ile or intragenic MTR ≤ 50^th^ %ile |
| **raredmg** (Rare damaging) | ≤ 0.005% | ≤ 0.025% | ≤ 0.005% | Missense | ≥ 0.25 | - |
| **raredmgmtr** (Rare damaging, MTR informed) | ≤ 0.005% | ≤ 0.025% | ≤ 0.005% | Missense | ≥ 0.25 | MTR ≤ 25^th^ %ile or  intragenic MTR ≤ 50^th^ %ile |
| **flexdmg** (Flexible MAF, damaging non-synonymous) | ≤ 0.1% (popmax) | ≤ 0.1% | ≤ 0.01% | Non-synonymous | ≥ 0.25 | - |
| **flexnonsyn** (Flexible MAF, all non-synonymous) | ≤ 0.1% (popmax) | ≤ 0.1% | ≤ 0.01% | Non-synonymous | - | - |
| **flexnonsynmtr** (Flexible MAF, non-synonymous, MTR informed) | ≤ 0.1% (popmax) | ≤ 0.1% | ≤ 0.01% | Non-synonymous | - | MTR ≤ 25^th^ %ile or  intragenic MTR ≤ 50^th^ %ile |
| **ptvraredmg** (PTV or rare damaging models combined) | PTV ≤ 0.1% (popmax) missense ≤ 0.005%  and ≤ 0.05% (popmax) | PTV ≤ 0.1% missense ≤ 0.025% | ≤ 0.01% | Non-synonymous | ≥ 0.25 | - |
| **rec** (Non-synonymous recessive) | ≤ 1% (popmax)  ≤ 10 homozygous calls | ≤ 1% | ≤ 0.1% | Non-synonymous | - | - |

* reflects the gnomAD global_raw MAF unless otherwise specified.
^ reflects the maximum proportion of UKB exome sequences permitted to either have ≤ 10-fold coverage at variant site or carry a low-confidence variant that did not meet one of the quality-control thresholds applied to collapsing analyses (see methods).
**Synonymous**: synonymous_variant
**PTV**: exon_loss_variant, frameshift_variant, start_lost, stop_gained, stop_lost, splice_acceptor_variant, splice_donor_variant, gene_fusion, bidirectional_gene_fusion, rare_amino_acid_variant, transcript_ablation
**Missense**: missense_variant_splice_region_variant, missense_variant
**Non-synonymous**: exon_loss_variant, frameshift_variant, start_lost, stop_gained, stop_lost, splice_acceptor_variant, splice_donor_variant, gene_fusion, bidirectional_gene_fusion, rare_amino_acid_variant, transcript_ablation, conservative_inframe_deletion, conservative_inframe_insertion ,disruptive_inframe_insertion, disruptive_inframe_deletion, missense_variant_splice_region_variant, missense_variant, protein_altering_variant

### Supplementary Table 4 – Collapsing analysis top hits

Provided as separate additional file

Associations from gene-level rare variant collapsing analysis where p<1x10^-8^. (Please note, our study-wide significance threshold is p<5 ×10^-9^). There are 1395 associations listed in total for binary traits of which 424 are significant and unique (i.e. counting associations that are significant in multiple models only once). There are 1652 associations listed in total for quantitative traits of which 487 are significant and unique.

### Supplementary Table 5 – Oncology aligned collapsing analyses

| **Gene** | **Category** | **Site(s)** |
| --- | --- | --- |
| *APC* | Solid | Digestive Organs |
| *ATM* | Solid | Digestive Organs |
| *BRCA1* | Solid | Breast, Female Genital Organs |
| *BRCA2* | Solid | Breast, Female Genital Organs, Male Genital Organs |
| *CDKN2A* | Solid | Skin |
| *CHEK2* | Solid | Breast |
| *MLH1* | Solid | Digestive Organs |
| *MSH6* | Solid | Digestive Organs, Female Genital Organs |
| *MUTYH* | Solid | Digestive Organs |
| *NF1* | Solid | Nervous System |
| *PALB2* | Solid | Breast |
| *RB1* | Solid | Eye |
| *ASXL1* | Haematological | Haematological |
| *DNMT3A* | Haematological | Haematological |
| *IGLL5* | Haematological | Haematological |
| *SRSF2* | Haematological | Haematological |
| *TET2* | Haematological | Haematological |

Summary of collapsing analysis results focusing on all genes achieving a p<1x10^-8^ in the Neoplasms Chapter. The full set of underlying oncology-aligned results can be found in **–Supplementary Table 6**. Although for a given gene there might be multiple correlated phenotypes and correlated collapsing models that achieve significance here for a given gen we summarise all the implicated sites. Secondary and benign neoplasms were excluded.

### Supplementary Table 6 - Oncology aligned collapsing analyses full

Provided as separate additional file

Gene-phenotype associations (p<1x10^-8^) with binary oncological traits in the collapsing analysis. These comprise both germline-driven hereditary solid tumours, and haematological malignancies even though the study was not designed to detect somatic variant-driven associations.

### Supplementary Table 7 – Control-enriched collapsing analyses

| **model** | **phenotype** | **Gene** | **Case Freq** | **Ctrl Freq** | **p-value** | **OR** | **OR LCI** | **OR UCI** | **OMIM** |
| --- | --- | --- | --- | --- | --- | --- | --- | --- | --- |
| flexdmg | Source of report of K80 (cholelithiasis) | *ABCG5* | 0.235% | 0.663% | 9.15E-09 | 0.3519 | 0.2325 | 0.5329 | Sitosterolemia; 210250 (3); Autosomal recessive |
| ptv | Union#E78#E78 Disorders of lipoprotein metabolism and other lipidaemias | *APOB* | 0.042% | 0.180% | 8.53E-12 | 0.2361 | 0.1436 | 0.3881 | Hypercholesterolemia; due to ligand-defective apo B; 144010 (3); Autosomal dominant\|Hypobetalipoproteinemia; 615558 (3); Autosomal recessive |
| ptv5pcnt | Union#J459#J45.9 Asthma\| unspecified | *IL33* | 0.692% | 1.177% | 9.44E-09 | 0.5853 | 0.4811 | 0.712 | . |
| ptv5pcnt | Union#E78#E78 Disorders of lipoprotein metabolism and other lipidaemias | *PCSK9* | 0.052% | 0.169% | 7.97E-09 | 0.3103 | 0.1972 | 0.4881 | Hypercholesterolemia; familial; 3; 603776 (3)\|{Low density lipoprotein cholesterol level QTL 1}; 603776 (3) |
| flexdmg | Union#M10#M10 Gout | *SLC22A12* | 0.061% | 0.477% | 4.56E-09 | 0.1264 | 0.0472 | 0.3383 | Hypouricemia; renal; 220150 (3); Autosomal recessive |

Summary of collapsing analysis results focusing on genes achieving a p<1x10^-8^ for control enrichment. Although for a given gene there might be multiple correlated phenotypes and correlated collapsing models that achieve significance here for a given gene we present the phenotype and collapsing model combination that achieved the lowest odds ratio. The major histocompatibility complex (MHC) region was excluded.

Supplementary Table 8: Comparison of significant gene-phenotype associations identified using collapsing and single variant (exWAS) analyses
The table provides the percentage of distinct gene-phenotypes associations (for both binary and
quantitative phenotypes) from the collapsing analysis that were also detected in the exWAS

|  | **Binary phenotypes** | **Quantitative phenotypes** |
| --- | --- | --- |
| **Total number of distinct gene-** **phenotype associations identified**  **in the collapsing analysis** | 424 | 487 |
| **Number of gene-phenotype associations from the collapsing analysis detected in the single-** **variant analysis i.e. exWAS (%)** | 70 (16.5%) | 311 (63.9%) |

[For this comparison, a significance threshold of p≤5x10^-9^ was used for both the collapsing analysis and the exWAS. Associations related to operation phenotypic codes (41200 and 41272) have been excluded for this comparison]

### Supplementary Table 9 – Non-intersecting PTV signals

Provided as separate additional file

Unique gene-trait pairs for statistically significant associations (p-value ≤ 5x10^-9^) discovered by two collapsing models that aggregate protein truncating variants with MAF < 0.1% and 5% ("ptv" and "ptv5pcnt", respectively). If the same gene-trait pair is found to be statistically significant (p-value ≤ 1x10^-8^) by single-variant tests (exWAS) using a single protein truncating variant for any of our exWAS models (genotypic, dominant, recessive), these associations are listed in column "exWAS". If any collapsing or exWAS associations do not reach our statistical significance thresholds, the equivalent cell is filled by "NA".

### Supplementary Table 10 – Null distribution

Provided as separate additional file

Binary_Syn: The 50 gene-phenotype associations (binary traits) from the synonymous negative control model of the collapsing analysis that have the lowest p values. Quant_Syn: The 50 gene-phenotype associations (quantitative traits) from the synonymous negative control model of the collapsing analysis that have the lowest p values. Binary_perms: The 50 gene-phenotype associations (binary traits) from the n-of-1 permutation based collapsing analysis that have the lowest p values. Quantit_perms: The 50 gene-phenotype associations (quantitative traits) from the n-of-1 permutation based collapsing analysis that have the lowest p values.

### Supplementary Table 11 – Gene Informativeness

Provided as separate additional file

Coverage statistics for each gene in CCDS release 22. Avg %10xCov Participant = percent of protein-coding sites covered with at least 10x coverage, averaged across all 177,882 UKB participants.

### Supplementary Table 12 – Cross-study 50k comparison

Provided as separate additional file

Unique gene-trait pairs for associations discovered by Regeneron, Helix and AstraZeneca using the first tranche of 50,000 exomes from UK Biobank. We only report associations with p-value<3.4x10^-10^, which is the significance threshold adopted by Helix publication. For the results from Helix and AstraZeneca, if the same gene-trait pair is found significant for different models (e.g. ptv versus coding for Helix), we used the pair with the highest statistical significance. For determining which traits were analysed by Regeneron, we used the 215 quantitative & 1023 binary traits listed in their Supplementary Table TraitsLists.xlsx.^3^ For Helix, we used the significant associations from the UK Biobank using European samples, found in Supplementary Data 2 from their manuscript, whereas all traits run by Helix are listed in their Supplementary Data 1.^4^
